# CRISPR-Cas-mediated tethering recruits the yeast *HMR* mating-type locus to the nuclear periphery but fails to silence gene expression

**DOI:** 10.1101/2021.04.30.442211

**Authors:** Emily R. Cliff, Robin L. Kirkpatrick, Daniel Cunningham-Bryant, Brianna Fernandez, Jesse G. Zalatan

## Abstract

To investigate the relationship between genome structure and function, we have developed a programmable CRISPR-Cas system for nuclear peripheral recruitment in yeast. We benchmarked this system at the *HMR* and *GAL2* loci, both well-characterized model systems for localization to the nuclear periphery. Using microscopy and gene silencing assays, we demonstrate that CRISPR-Cas-mediated tethering can recruit the *HMR* locus but does not silence reporter gene expression. A previously reported Gal4-mediated tethering system does silence gene expression, and we demonstrate that the silencing phenotype has an unexpected dependence on the structure of the protein tether. The CRISPR-Cas system was unable to recruit *GAL2* to the nuclear periphery. Our results reveal potential challenges for synthetic genome structure perturbations and suggest that distinct functional effects can arise from subtle structural differences in how genes are recruited to the periphery.

## Introduction

An emerging body of data suggests that the 3D spatial organization of the genome plays an important role in eukaryotic gene regulation.^1–4^ For example, genes positioned near the nuclear periphery tend to be repressed, and genes positioned in the nuclear interior tend to be active.^3–5^ Further support for the possibility that genome organization plays an important regulatory role comes from the observation that genes can be dynamically repositioned upon activation or in response to extracellular signals.^6–9^ In contrast, genome structures can undergo major perturbations with only modest effect on the transcriptome when cohesion-mediated loops are disrupted in human cells.^10^ Therefore, tools are needed which allow us to systematically probe the biological function of genome structure. To this end, we have developed a programmable CRISPR-Cas system to relocalize genes to the nuclear periphery and prototyped the system in yeast.

Prior methods to reposition genes have fused well characterized DNA binding domains (DBD), such as Gal4, to a recruitment domain protein that directs the tethered gene to specific sites in the nucleus.^7,11–17^ In these studies, repositioning genes sometimes, but not always, leads to predictable changes in gene expression. The number of sites that have been studied with this approach has been relatively limited, in part because DBDs typically recognize specific DNA sequences and these motifs must be engineered into each genomic site of interest.

To address this challenge, CRISPR-Cas tethering systems have been developed to target and spatially reposition genomic sites within the nucleus. ^18–23^ Because CRISPR-Cas targeting is programmable, such systems enable recruitment of endogenous genes and bypass the need for site specific gene modification of the recruitment target site. Several of these systems enable recruitment of genomic sites to the nuclear periphery. In human cells, dCas9 fusion to a chemically-inducible dimerization domain allowed inducible recruitment to the nuclear envelope and other subnuclear sites. Reporter and endogenous gene expression could be perturbed by nuclear repositioning with this system, and localization of telomeres to the nuclear periphery resulted in cellular toxicity.^18^ In a separate study, direct fusion of dCas9 to the lamin protein Lap2β also enabled peripheral recruitment.^19^ In yeast cells, dCas9 fusion to a cohesin domain could target a dockerin fused to a nuclear membrane protein. This system successfully recruited multiple endogenous loci to the nuclear periphery and was able to affect plasmid segregation to daughter cells.^20^ These findings suggest that CRISPR gene relocalization systems could be useful to discover relationships between gene positioning and cellular behavior.

In parallel to the methods described above, we developed an alternative CRISPR-Cas repositioning system in yeast and tested it at the *HMR* locus, a well-characterized model system for position-dependent gene silencing. Gal4-mediated recruitment of *HMR* to the nuclear periphery can rescue gene silencing defects, ^11^ and we assessed whether a CRISPR-Cas-mediated recruitment strategy would have the same functional effects. Using a nuclear membrane protein as the recruitment domain, we targeted either the CRISPR-Cas system or the Gal4 system to the *HMR* locus and measured nuclear peripheral relocalization by microscopy. We found that both systems produce similar, significant levels of recruitment. Next, we compared the ability of CRISPR-Cas and Gal4 recruitment systems to modulate gene expression. We found that only the Gal4 system was able to silence the *HMR* locus, but this system was unexpectedly sensitive to the structure of the Gal4-membrane protein fusion. Inserting a protein spacer between Gal4 and the nuclear membrane protein maintains recruitment but abrogates silencing. This result suggests that although alternative tethering strategies can be used to recruit genes to the periphery, silencing and other functional effects may depend on the precise structural orientation. We also tested the CRISPR-Cas recruitment system at the *GAL2* locus, which is a model for dynamic repositioning in response to external stimuli.^6,24–26^ We were unable to recruit *GAL2* to the nuclear periphery with the CRISPR-Cas system, suggesting that not all endogenous target sites can be synthetically relocalized to the nuclear periphery.

## Results

### Gal4 can recruit the HMR locus to the nuclear periphery and silence gene expression

The yeast *HMR* locus provided one of the earliest examples of a functional effect from synthetic gene repositioning, making it an ideal model system to prototype a new repositioning system. *HMR* contains a backup copy of yeast mating type sequences and is natively silenced.^27^ Deletion of *HMR* regulatory regions results in locus de-repression,^28^ and silencing can be restored by synthetically recruiting a Gal4 DNA binding domain fused to a transcriptional repressor.^29^ Gene silencing can also be restored by Gal4-mediated tethering of nuclear membrane proteins to *HMR*. This effect is thought to be due to repositioning of *HMR* to a nuclear peripheral location with high concentrations of silencing factors.^11^

To compare the ability of Gal4 or dCas9 to recruit *HMR* to the nuclear periphery in yeast, we constructed an *HMR* silencing reporter in *S. cerevisiae*. Following previous designs, ^11,29^ we replaced the endogenous copy of *HMR* with a mutant in the *HMR-E* regulatory region to de-repress the locus. The mutant cassette also includes a 2xUAS_G_ site upstream of a Trp1 reporter (Aeb::2xUAS_G_ *hmr*::*Trp1*) (see Methods). We fused the Gal4 DBD to the nuclear membrane protein Yif1 (Fig 1) and confirmed that Gal4DBD-Yif1 expression silences the Trp1 reporter gene in a cell-spotting growth assay, as described previously (see below & Fig 3).^11^ To determine if Gal4DBD-Yif1 physically repositions *HMR* to the nuclear periphery, which was not previously assessed, we further engineered the silencing reporter strain with a tetO array 2.4 kb downstream of the UAS_G_ site (Fig 2A).^30^ Expression of tetR-GFP allows visualization of the *HMR* locus. We expressed the mCherry-Heh2 fusion protein to label the nuclear membrane,^31,32^ and used confocal microscopy to measure the position of the tetO array relative to the nuclear rim (Fig 2B). We observed a significant change in *HMR* peripheral localization when Gal4DBD-Yif1 was expressed compared to Gal4DBD alone, increasing from 39% to 51% (Fig 2C).

**Figure 1.**
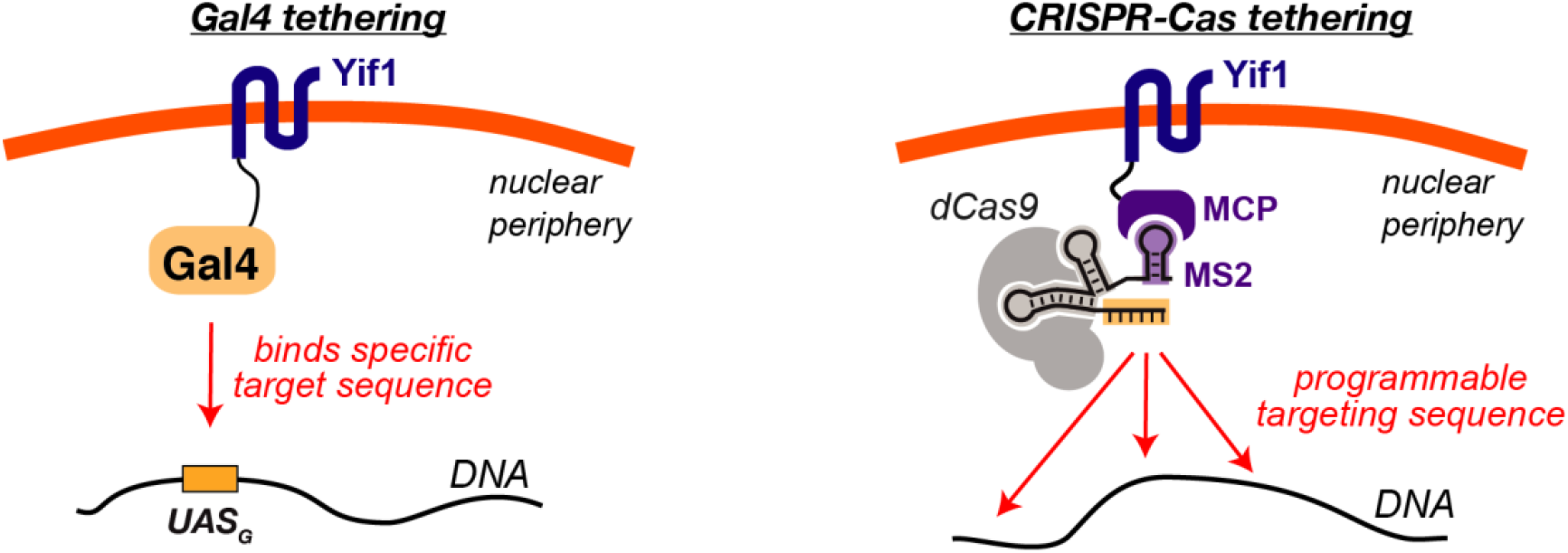
Peripheral tethering with nuclear membrane fusions. Genomic sites can be physically repositioned by fusing a DNA binding domain to a membrane protein. The DNA binding domain can be a protein like Gal4, which binds a specific DNA target, or a CRISPR-Cas complex that can be programmed to different target sites. The CRISPR-Cas complex can be linked to a membrane protein via a scaffold RNA (scRNA), a modified gRNA that includes an MS2 RNA hairpin to recruit the MS2 coat protein (MCP) fused to the membrane protein Yif1.

**Figure 2.**
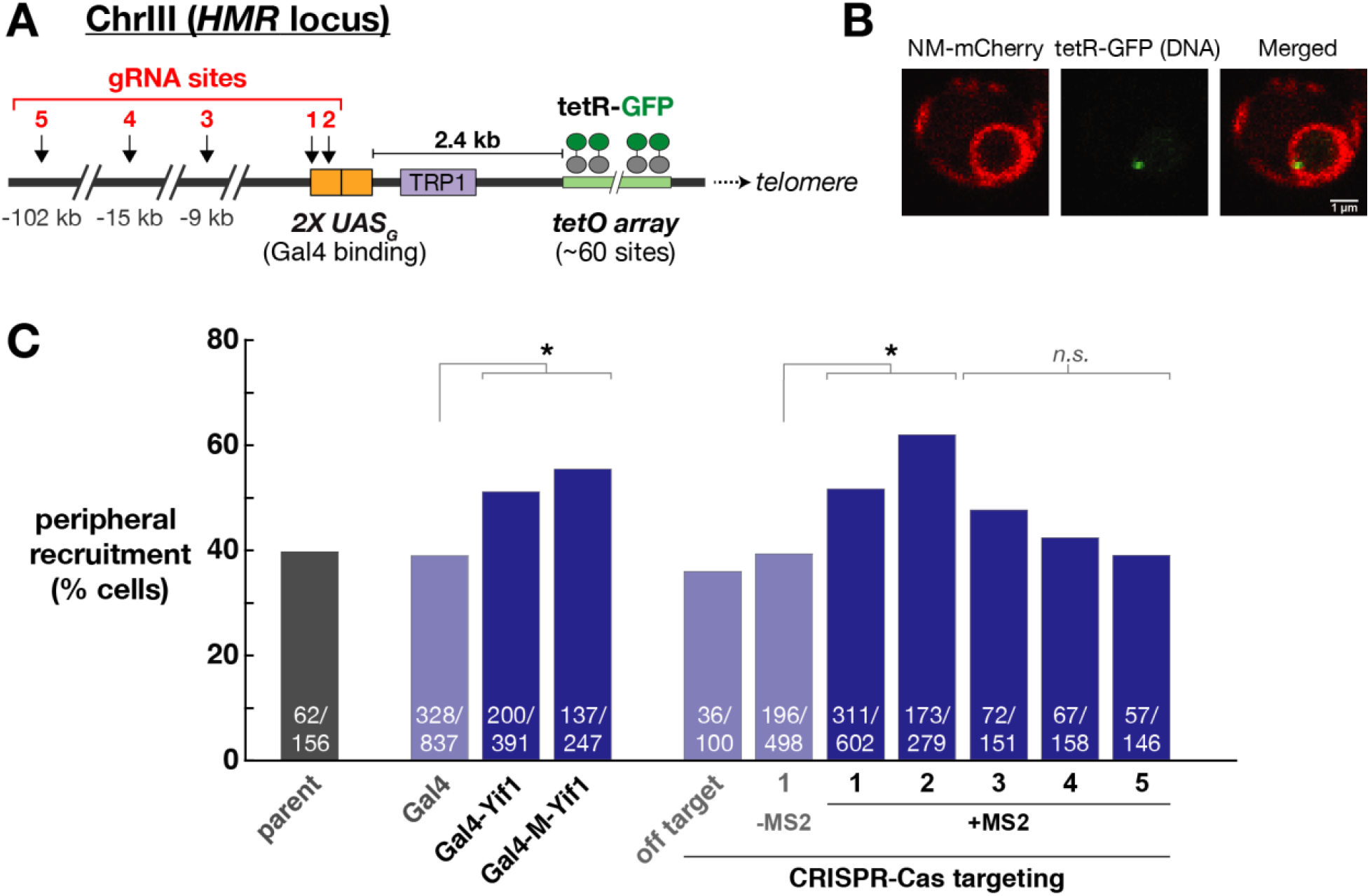
Visualization of *HMR* peripheral recruitment by microscopy. (A) The *HMR* locus was engineered with a 2X UAS_G_ site, a Trp1 reporter, and a tetO array. See Supporting Information for a complete annotated sequence. (B) Representative confocal microscopy images show the nuclear envelope (red), defined by an mCherry-Heh2 fusion protein, and the *HMR* locus (green), defined by tetR-GFP that binds the tetO array. (C) Peripheral recruitment was scored as described in the methods for yeast strains with and without recruitment systems. The parent yeast strain (yRK119) is indistinguishable from negative control strains with an off-target scRNA (OT, see Table S1) or a gRNA lacking the MS2 recruitment hairpins (–MS2). At least 100 cells were measured for each strain. Exact values (recruited/total) are shown in white text with each bar. Statistical significance for a significant change in localization relative to a corresponding negative control was evaluated using a 2-tailed chi-squared test (*p* value ≤ 0.05, indicated by *). The *p* values for targets 1-5, relative to the –MS2 control, are <0.0001, <0.0001, 0.08, 0.56, and 0.94, respectively. Gal4_DBD_-Yif1 and Gal4_DBD_-M-Yif1 (containing MBP between Gal4_DBD_ and Yif1) also show significant peripheral recruitment relative to a strain containing only Gal4_DBD_ (both *p* values <0.0001).

**Figure 3.**
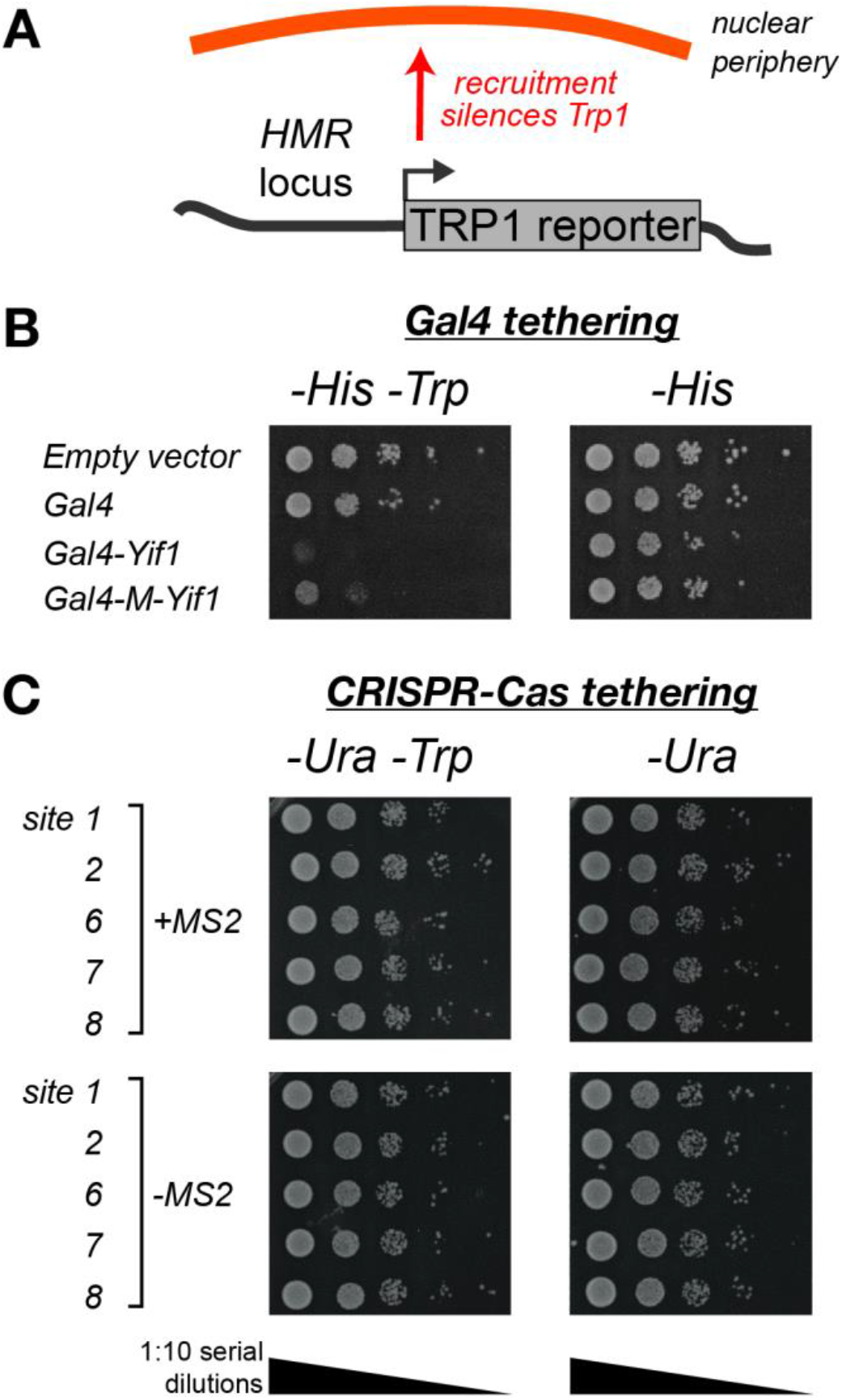
Trp1 reporter silencing assay. (A) Peripheral recruitment of the *HMR* locus silences a Trp1 reporter gene, leading to a growth defect on media lacking Trp. See Fig 2A for a complete schematic of the *HMR* locus and the Trp1 reporter. (B) Gal4DBD-Yif1 expression results in a growth defect on –His –Trp media, as previously reported.^11^ Cells expressing Gal4_DBD_-M-Yif1, which has MBP inserted between Gal4_DBD_ and Yif1, produce a partial silencing phenotype. The parent strain is yRK036 and Gal4 constructs were delivered on p423 (His selection). (C) A CRISPR-Cas complex that recruits *HMR* to the periphery does not silence the Trp1 reporter gene. There is no detectable growth defect on –Ura –Trp media. See Table S1 for guide RNA sequences and locations. The parent strain is yRK045, and 2x MS2 scRNA or –MS2 sgRNA (negative control) constructs were delivered on pRS316 (Ura selection). Images in (B) and (C) are representative of three independent experiments (biological replicates).

### CRISPR-Cas can recruit the HMR locus to the nuclear periphery

To determine if the CRISPR-Cas system can relocalize the *HMR* locus, we physically linked the CRISPR-Cas complex to Yif1, the same nuclear membrane protein used in the Gal4 tethering strategy. We linked Yif1 to the CRISPR-Cas complex through a scaffold RNA (scRNA), which is an sgRNA engineered with additional hairpin motifs to recruit effectors fused to an RNA binding protein (Fig 1).^33–36^ In this strategy, Yif1 is fused to the MS2 coat protein (MCP), which binds as a dimer to an MS2 hairpin on the scRNA (Fig S1). To recruit dCas9 to the *HMR* locus, we targeted sites adjacent to or overlapping the UAS_G_ site with scRNA constructs containing two MS2 RNA hairpins. We transformed individual scRNA constructs separately into the reporter yeast strain expressing dCas9 and MCP-Yif1. Using confocal microscopy, we observed a significant increase in peripheral localization with the scRNA-containing strains (Fig 2). At both UAS_G_-adjacent sites initially tested (sites 1 and 2), a single scRNA was sufficient to reposition the *HMR* locus. Guide RNAs with an off-target (OT) sequence or lacking the MS2 hairpins (–MS2) gave no significant peripheral recruitment, as expected. We also tested a direct dCas9-Yif1 fusion protein and observed similar recruitment effects (Fig S2).

Both dCas9 and Gal4 recruitment strategies resulted in similar recruitment phenotypes (Fig 2C). For both the CRISPR-Cas and the Gal4 recruitment strategies, the 50-60% peripheral localization that we observe is comparable to the 50-80% range of values observed for endogenous and heterologous peripheral recruitment reported in the literature.^7,32,37,38^ The background peripheral localization for the unrecruited *HMR* silencing reporter was ~40%, which is modestly larger than the 30% peripheral localization values reported for unrecruited genes in other systems. This larger background for the *HMR* reporter may be due to its proximity to the ChrIII telomere, as telomeres are endogenously localized to the periphery in yeast.^39,40^

In addition to targeting sites relatively close the tetO array (sites 1 and 2, within 2.4 kb), we also tested scRNA target sites at increasing distances. We designed MS2 scRNAs to target additional sites 9 kb, 15 kb, and 102 kb from the tetO array. For each of these sites, we did not detect statistically significant increases in peripheral localization (Fig 2C). These data suggest that the target site and microscopy reporter need to be in close physical proximity to observe gene relocalization. How these distances in base pairs translate to physical distances is uncertain. Using a previously described yeast chromatin polymer model with a linear mass density of 144 nm/bp, ^41^ we can roughly estimate that these sites are 57 nm, 97 nm, and 451 nm respectively from the tetO array, but we lack any direct measurements of the distances to *HMR* for these specific sites. Regardless of the precise distance relationship, however, our data suggest that a CRISPR-Cas-based gene recruitment system can localize nearby genomic regions to the periphery using a single scRNA targeting a unique site in the genome. As the distance to the target site increases, the genomic locus of interest is less likely to be repositioned.

### Peripheral recruitment is not sufficient to silence reporter gene expression

To determine if CRISPR-Cas-mediated *HMR* recruitment produced the same silencing effect on gene expression as observed with Gal4-Yif1, we assessed Trp1 reporter gene expression using the cell-spotting growth assay. Although CRISPR-Cas-mediated tethering to Yif1 and Gal4_DBD_-Yif1were indistinguishable by microscopy (Fig 2C), there was no detectable silencing at *HMR* with the CRISPR-Cas system (Fig 3). In addition to the two scRNA target sites used in microscopy assays, we tested three additional nearby target sites but did not observe Trp silencing with any of these sites (Fig 3C). There are no immediately obvious distinguishing features of the CRISPR-Cas and Gal4 systems that could explain their distinct behaviors. The affinities of the recruitment interactions are all relatively similar, in the range of ~10^−9^ M, for Gal4 dimer binding to the UAS_G_,^42^ dCas9-gRNA binding to cognate DNA, ^43,44^ and the MS2 RNA hairpin for the MCP dimer (the MCPV29IΔFG variant).^45^ Gal4 binds the UAS_G_ site as a dimer, so a 2X UAS_G_ target recruits four copies of Gal4-Yif1. MCP is also a functional dimer, and a single 2x MS2 scRNA recruits four copies of MCP-Yif1 (Fig S1).

Gal4DBD-Yif1-mediated silencing at *HMR* is known to require the presence of endogenous cis-regulatory sites,^11^ and it is possible that the precise structural arrangement of the Gal4_DBD_-Yif1 fusion protein relative to these sites or other associated regulatory factors might be important for silencing. To test this possibility, we inserted maltose binding protein (MBP) between Gal4_DBD_ and Yif1 (Gal4_DBD_-M-Yif1). MBP is typically used as a protein affinity tag for purification. In this context we expect MBP to be an inert spacer that extends the distance between Gal4DBD and Yif1 by ~40 Å (estimated from the crystal structure of MBP).^46^ By microscopy, Gal4DBD-M-Yif1 resulted in peripheral localization that was indistinguishable from Gal4_DBD_-Yif1 or CRISPR-Cas recruitment (Fig 2C). Unlike Gal4_DBD_-Yif1, however, the Gal4_DBD_-M-Yif1 construct produced only a partial silencing phenotype (Fig 3B). This observation suggests that, at least at the *HMR* locus in yeast, peripheral gene silencing may depend on the precise structure of the recruitment machinery.

### The CRISPR-Cas system does not recruit the GAL2 locus to the nuclear periphery

To determine if CRISPR-Cas-mediated recruitment is effective at other sites in the genome, we targeted the *GAL2* locus. In response to galactose, yeast cells localize *GAL2* to the nuclear pore complex and activate *GAL2* expression by ~20-fold.^6,24,47^ This behavior indicates that it is possible to reposition *GAL2* and provides a useful positive control for comparison to synthetic recruitment strategies. We constructed a reporter strain with a tetO array at the *GAL2* locus to visualize its position and confirmed that galactose induction recruits *GAL2* to the periphery (Fig 4A, B). The extent of peripheral localization increased from 32% in glucose to 47% in galactose, comparable to previously reported results.^24^ When we used a Yif1-tethered scRNA to target the CRISPR-Cas system to *GAL2*, however, we did not detect significant repositioning to the nuclear periphery (Fig 4C). We also tested a system with simultaneous expression of four scRNAs targeting adjacent sites at *GAL2*, but again observed no significant repositioning (Fig 4C). Thus, although our CRISPR-Cas recruitment system was effective at peripheral tethering of *HMR*, we were unable to detect repositioning at a different site in the yeast genome. This behavior is in contrast with an alternative recruitment system, CRISPR-PIN, which was able to effectively recruit multiple distinct endogenous loci in yeast.^20^ Our system uses a different recruitment domain, Yif1, but this protein has been used as a LexA-Yif1 fusion to recruit other yeast genomic sites to the nuclear periphery.^12^ In principle, these precedents suggest that a CRISPR-Cas system with Yif1 should be effective at other genomic sites besides *HMR*, but our results indicate that the system may not be effective at arbitrary genomic loci. Synthetic recruitment at some sites could be limited by pre-existing genome structure that is resistant to repositioning, weak binding of the CRISPR-Cas complex due to ineffective gRNA target sequence, or the presence of inaccessible chromatin.^48,49^

**Figure 4.**
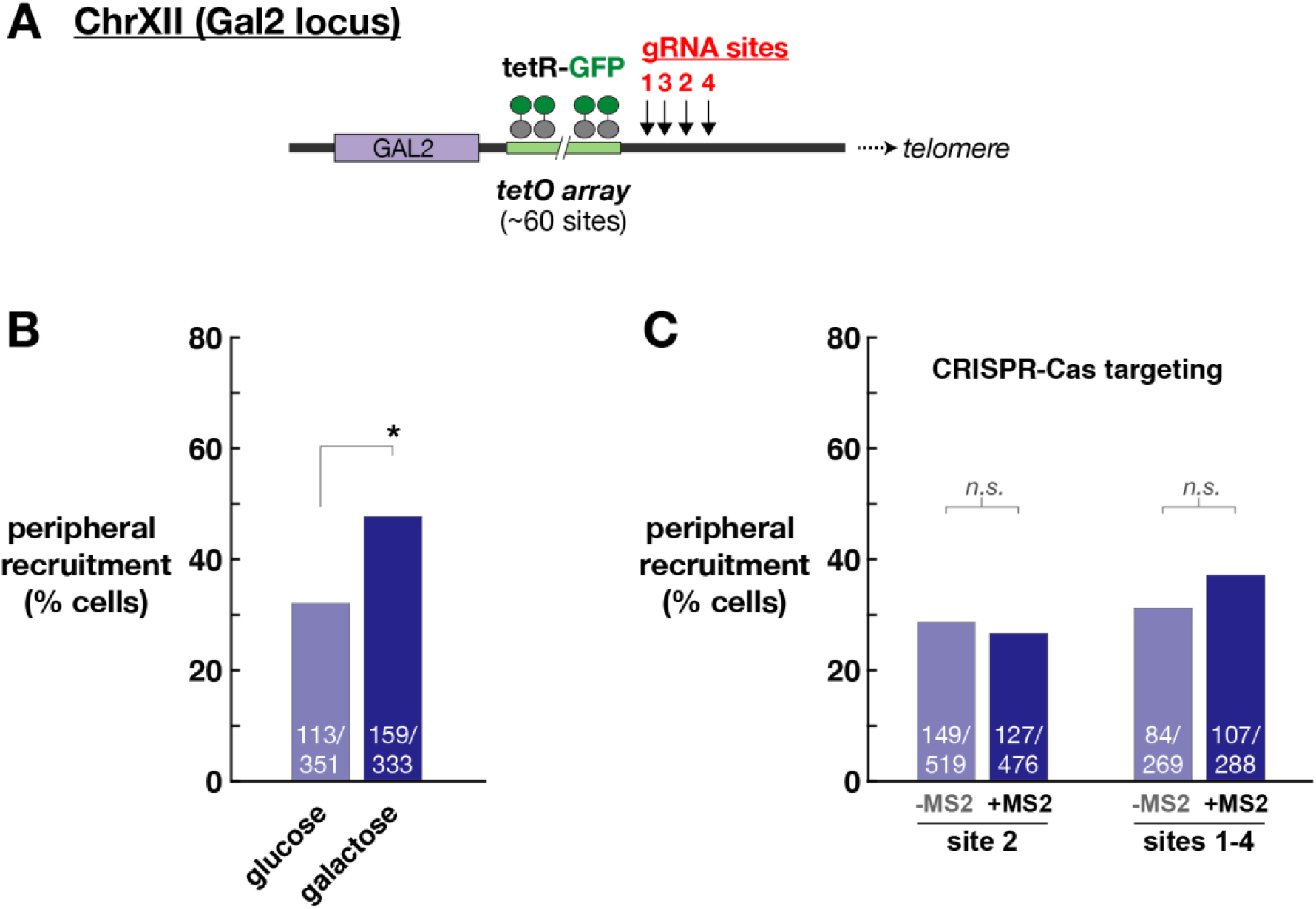
Visualization of *GAL2* peripheral recruitment by microscopy. (A) The *GAL2* locus was engineered with a tetO array. See Supporting Information for a complete annotated sequence. (B) Growth in galactose recruits *GAL2* to the nuclear periphery. (C) *GAL2* was targeted with the CRISPR-Cas tethering system with one (site 2) or four (sites 1-4) 2X MS2 scRNAs. gRNAs lacking MS2 (-MS2) were used as negative controls. No significant recruitment was detected. For (B) and (C), at least 100 cells were measured for each strain. Exact values (recruited/total) are shown in white text with each bar. Statistical significance for a change in localization relative to the corresponding negative control was evaluated using a chi-squared test (indicated by *). The *p* value for growth in galactose relative to glucose is <0.0001. The *p*-values for CRISPR-Cas tethering attempts were 0.52 (site 2) and 0.17 (all four sites).

## Discussion

In this study, we developed a programmable CRISPR-Cas system for recruiting genes to the nuclear periphery in yeast. Our initial experiments demonstrated that Gal4 and CRISPR-Cas tethering systems could recruit the *HMR* locus to the nuclear periphery, and the recruitment effects were indistinguishable by microscopy. However, the CRISPR-Cas gene tethering system was unable to reproduce the gene silencing effects of the Gal4-mediated system. Further, modifying the Gal4 system with an MBP protein between the Gal4 DBD and the Yif1 recruitment domain maintained the recruitment effect but substantially weakened the gene silencing phenotype. These results suggest that, while these systems produce indistinguishable peripheral recruitment at the resolution of our microscopy assay, there are unresolved structural differences that lead to distinct functional effects.

It is well established that synthetic peripheral recruitment can repress genes in yeast and metazoan cells.^16,50^ Although the underlying molecular mechanism remains unclear, several plausible models have been suggested. One possibility is that peripheral recruitment brings genes into close proximity to silencing factors that are already localized at the periphery. ^16,50^ Alternatively, peripheral recruitment could sequester genes away from compartments in the nuclear interior where transcriptional machinery is localized and active. ^16^ Adding to the uncertainty is the observation by several groups that peripheral recruitment does not always lead to silencing.^14,51^

In yeast cells, several experiments support the idea that localized silencing factors are important for peripheral silencing. Localized Sir proteins are necessary for synthetic peripheral silencing in yeast, and disrupting Sir protein localization allows silencing at internal genes.^11,52^ Further, tethering and silencing can be decoupled by mutations that prevent formation of the Sir complex.^12^ In the *HMR* reporter used in our experiments, a cis-regulatory DNA sequence (the A site) is necessary for silencing by synthetic peripheral recruitment.^11,29^ Peripheral recruitment of *HMR* may bring the locus to a peripheral location where high concentrations of Sir proteins are able to bind at this site and silence the reporter.

Both models to explain peripheral silencing, localized silencing factors or sequestration from active compartments, predict that repression should be independent of the exact method used for peripheral recruitment. Instead, we find that perturbing the structure of the Gal4-Yif1 tether, or switching to a CRISPR-Cas tethering system, maintains tethering but fails to silence gene expression. There may be subtle differences in precise positioning or orientation that lead to distinct functional effects from the different tethering systems. Alternatively, recent work has highlighted a potential role for phase separation in the formation of heterochromatin, ^53^ and it is possible that some tethering systems could interfere with physical partitioning into a silenced region. In either case, our results highlight an important reminder for synthetic biology that systems which appear to be modular and have similar physical properties do not necessarily have the same functional effects.

## Methods

### Yeast Strain Construction and Manipulation

Yeast (*S. cerevisiae*) transformations were performed with the standard lithium acetate method. The parent haploid yeast strain for reporter gene experiments was SO992 (W303;*MAT***a** *ura3 leu2 trp1 his3*). Complete descriptions of all yeast strains generated in this work are provided in Table S2, and descriptions of the plasmids are in Table S3. Complete sequences for guide RNAs, effector proteins, and reporter genes are providing in the Supporting Information. dCas9 and MCP fusion proteins were expressed as described previously from constructs integrated in single copy into the yeast genome.^35^ Yif1 was cloned from yeast genomic DNA. Yif1 fusion proteins used Yif1(55-314)^11^ fused to the C-terminus of Gal4DBD, dCas9, or MCP. Guide RNA constructs were expressed as described previously from the pRS316 CEN/ARS plasmid (*ura3* marker) with the SNR52 promoter and SUP4 terminator.^35^ For simultaneous expression of four unique guide RNAs, multiple guide cassettes with independent SNR52 promoters and SUP4 terminators were expressed from a single pRS316 plasmid.^54^ All Gal4_DBD_ constructs and derivatives were expressed from the p423 2µ plasmid. The *HMR* Trp1 reporter strain (Aeb::2xUAS_G_ *hmr*::*Trp1*) at the endogenous *HMR* locus was constructed by transforming a linear DNA fragment (derived from plasmid pRK105) containing *HMR*-Aeb_2xUAS_G__Trp1 with >280 bases of flanking homology and selecting on SD –Trp plates. Integration of the full reporter cassette was verified by colony PCR. TetO arrays were integrated at ChrIII (*HMR*) and ChrXII (*GAL2*) using either pSR8 (*his3* marker) or pSR14 (*leu2* marker) (gifts from Susan Gasser), using previously described methods. ^30^ Based on plasmid sizes, we estimate that our TetO arrays contained ~ 60-80 tetO repeats. The tetR-GFP and mCherry-Heh2138-378 fusion proteins were integrated in single copy in the yeast genome. The expression cassette for the tetR-GFP protein was derived from pGVH29 (gift from Susan Gasser).^41^ The mCherry-Heh2138-378 construct was designed following previous reports. ^31,32^

### Trp1 Silencing Assay

After transformations, yeast strains were grown overnight at 30 °C on selective plates (SD–Ura or SD –His as appropriate). Patches were diluted to OD_600_ 0.2 in selective media lacking Trp and serially diluted 1:10 to result in the following set of serial dilutions: N/A, 1:10, 1:100, 1:1000, 1:10000. 10 μL of each dilution was spotted on selective SD plates with or without Trp. Plates were incubated at 30 °C and evaluated after 2 days.

### Yeast Microscopy

After transformations, yeast strains were grown overnight at 30 °C on selective plates (SD–Ura or SD –His) or YPD (parent strain). Cells were resuspended in YPD at a starting OD_600_ ~0.15 and grown to OD600 0.3-0.5. 5 mL cultures were pelleted, washed in SD complete media, and resuspended in 20 µL SD complete.

For galactose-induced repositioning of *GAL2*, cells were grown overnight in YPRaf (yeast peptone with 2% raffinose). Cells were resuspended in either YPD (2% glucose) or YPGal (2% galactose) at a starting OD600 ~0.15 and grown to OD600 0.3-0.5. 5 mL cultures were then pelleted, washed in either SD (2% glucose) or SGal (2% galactose) as appropriate and resuspended in 20 µL of the same media.

For microscopy, 10 µL of resuspended cells were pipetted onto agarose pads in 13 x 1 mm silicone isolator wells (Electron Microscopy Sciences) and covered with a No. 1.5 coverslip. Imaging was performed using a Leica TCS SP5 II laser scanning confocal microscope with a 63x oil immersion objective. The pixel size was 90.1 nm and the z-step size was 0.21 µm. The optical thickness of each slice is 0.98 µm. For each cell, the tetR-GFP spot was assigned to a particular z-plane based on its maximum intensity. In that z-plane, we defined the nuclear periphery as the pixel corresponding to the center of the Heh2-mCherry peak along the radial axis. GFP spots were classified as “peripheral” if the center of spot was within two pixels of the nuclear periphery (i.e. separated by no more than one 90.1 nm pixel). Cells in which the GFP spot was assigned to the bottom or top slice of the nucleus were excluded from analysis. ^55^

## Acknowledgements

The authors thank Susan Gasser, Sue Biggins, Dan Gottschling, Joshua Vaughan, Tyler Chozinski, Dustin Maly, Jack Rose, Jay Shendure, Bill Noble, David Shechner, James Carothers, Jason Brickner, Wendell Lim, Geeta Narlikar, Wai Chan, the Biology Imaging Facility at the University of Washington, and members of the Zalatan group for technical assistance, advice, and helpful discussions. This work was supported by a Career Award at the Scientific Interface from the Burroughs Wellcome Fund (J.G.Z.), a Genome Sciences NIH Training Grant T32 HG00035 (R.L.K.), and NIH R35 GM124773 (J.G.Z.).

## Author Contributions

R.L.K., E.R.C., D.C.B., and J.G.Z. designed experiments. R.L.K., E.R.C., D.C.B., and B.F. performed experiments. R.L.K., E.R.C., and J.G.Z. wrote the manuscript.

## Notes

The authors declare no competing financial interest.

## Notes

### Competing Interest Statement

The authors have declared no competing interest.

## References

(1) Avşaroğlu, B., Bronk, G., Li, K., Haber, J. E., and Kondev, J. (2016) Chromosome-refolding model of mating-type switching in yeast. Proc. Natl. Acad. Sci. USA 113, E6929–E6938.

(2) Lieberman-Aiden, E., van Berkum, N. L., Williams, L., Imakaev, M., Ragoczy, T., Telling, A., Amit, I., Lajoie, B. R., Sabo, P. J., Dorschner, M. O., Sandstrom, R., Bernstein, B., Bender, M. A., Groudine, M., Gnirke, A., Stamatoyannopoulos, J., Mirny, L. A., Lander, E. S., and Dekker, J. (2009) Comprehensive mapping of long-range interactions reveals folding principles of the human genome. Science 326, 289–293.

(3) Egecioglu, D., and Brickner, J. H. (2011) Gene positioning and expression. Curr. Opin. Cell Biol. 23, 338–345.

(4) Ramani, V., Shendure, J., and Duan, Z. (2016) Understanding spatial genome organization: methods and insights. Genomics Proteomics Bioinformatics 14, 7–20.

(5) Misteli, T. (2020) The Self-Organizing Genome: Principles of Genome Architecture and Function. Cell 183, 28–45.

(6) Casolari, J. M., Brown, C. R., Komili, S., West, J., Hieronymus, H., and Silver, P. A. (2004) Genome-wide localization of the nuclear transport machinery couples transcriptional status and nuclear organization. Cell 117, 427–439.

(7) Brickner, J. H., and Walter, P. (2004) Gene recruitment of the activated INO1 locus to the nuclear membrane. PLoS Biol. 2, e342.

(8) Randise-Hinchliff, C., and Brickner, J. H. (2016) Transcription factors dynamically control the spatial organization of the yeast genome. Nucleus 7, 369–374.

(9) Kim, S., Liachko, I., Brickner, D. G., Cook, K., Noble, W. S., Brickner, J. H., Shendure, J., and Dunham, M. J. (2017) The dynamic three-dimensional organization of the diploid yeast genome. eLife 6.

(10) Rao, S. S. P., Huang, S.-C., Hilaire, B. G. S., Engreitz, J. M., Perez, E. M., Kieffer-Kwon, K.-R., Sanborn, A. L., Johnstone, S. E., Bascom, G. D., Bochkov, I. D., Huang, X., Shamim, M. S., Shin, J., Turner, D., Ye, Z., Omer, A. D., Robinson, J. T., Schlick, T., Bernstein, B. E., Casellas, R., Lander, E. S., and Aiden, E. L. (2017) Cohesin Loss Eliminates All Loop Domains. Cell 171, 305–309.e24.

(11) Andrulis, E. D., Neiman, A. M., Zappulla, D. C., and Sternglanz, R. (1998) Perinuclear localization of chromatin facilitate s transcriptional silencing. Nature 394, 592–595.

(12) Taddei, A., Hediger, F., Neumann, F. R., Bauer, C., and Gasser, S. M. (2004) Separation of silencing from perinuclear anchoring functions in yeast Ku80, Sir4 and Esc1 proteins. EMBO J. 23, 1301–1312.

(13) Menon, B. B., Sarma, N. J., Pasula, S., Deminoff, S. J., Willis, K. A., Barbara, K. E., Andrews, B., and Santangelo, G. M. (2005) Reverse recruitment: the Nup84 nuclear pore subcomplex mediates Rap1/Gcr1/Gcr2 transcriptional activation. Proc. Natl. Acad. Sci. USA 102, 5749–5754.

(14) Finlan, L. E., Sproul, D., Thomson, I., Boyle, S., Kerr, E., Perry, P., Ylstra, B., Chubb, J. R., and Bickmore, W. A. (2008) Recruitment to the nuclear periphery can alter expression of genes in human cells. PLoS Genet. 4, e1000039.

(15) Reddy, K. L., Zullo, J. M., Bertolino, E., and Singh, H. (2008) Transcriptional repression mediated by repositioning of genes to the nuclear lamina. Nature 452, 243–247.

(16) van Steensel, B., and Belmont, A. S. (2017) Lamina-associated domains: Links with chromosome architecture, heterochromatin, and gene repression. Cell 169, 780–791.

(17) Wang, H., Han, M., and Qi, L. S. (2021) Engineering 3D genome organization. Nat. Rev. Genet.

(18) Wang, H., Xu, X., Nguyen, C. M., Liu, Y., Gao, Y., Lin, X., Daley, T., Kipniss, N. H., La Russa, M., and Qi, L. S. (2018) CRISPR-mediated programmable 3D genome positioning and nuclear organization. Cell 175, 1405–1417.e14.

(19) See, K., Kiseleva, A. A., Smith, C. L., Liu, F., Li, J., Poleshko, A., and Epstein, J. A. (2020) Histone methyltransferase activity programs nuclear peripheral genome positioning. Dev. Biol. 466, 90–98.

(20) Lin, J.-L., Ekas, H., Deaner, M., and Alper, H. S. (2019) CRISPR-PIN: Modifying gene position in the nucleus via dCas9-mediated tethering. Synth Syst Biotechnol 4, 73–78.

(21) Morgan, S. L., Mariano, N. C., Bermudez, A., Arruda, N. L., Wu, F., Luo, Y., Shankar, G., Jia, L., Chen, H., Hu, J.-F., Hoffman, A. R., Huang, C.-C., Pitteri, S. J., and Wang, K. C. (2017) Manipulation of nuclear architecture through CRISPR-mediated chromosomal looping. Nat. Commun. 8, 15993.

(22) Hao, N., Shearwin, K. E., and Dodd, I. B. (2017) Programmable DNA looping using engineered bivalent dCas9 complexes. Nat. Commun. 8, 1628.

(23) Kim, J.-H., Rege, M., Valeri, J., Dunagin, M. C., Metzger, A., Titus, K. R., Gilgenast, T. G., Gong, W., Beagan, J. A., Raj, A., and Phillips-Cremins, J. E. (2019) LADL: light-activated dynamic looping for endogenous gene expression control. Nat. Methods 16, 633–639.

(24) Dieppois, G., Iglesias, N., and Stutz, F. (2006) Cotranscriptional recruitment to the mRNA export receptor Mex67p contributes to nuclear pore anchoring of activated genes. Mol. Cell. Biol. 26, 7858–7870.

(25) Sood, V., and Brickner, J. H. (2014) Nuclear pore interactions with the genome. Curr. Opin. Genet. Dev. 25, 43–49.

(26) Sood, V., Cajigas, I., D’Urso, A., Light, W. H., and Brickner, J. H. (2017) Epigenetic Transcriptional Memory of GAL Genes Depends on Growth in Glucose and the Tup1 Transcription Factor in Saccharomyces cerevisiae. Genetics 206, 1895–1907.

(27) Haber, J. E. (2012) Mating-type genes and MAT switching in Saccharomyces cerevisiae. Genetics 191, 33–64.

(28) Brand, A. H., Micklem, G., and Nasmyth, K. (1987) A yeast silencer contains sequences that can promote autonomous plasmid replication and transcriptional activation. Cell 51, 709– 719.

(29) Chien, C. T., Buck, S., Sternglanz, R., and Shore, D. (1993) Targeting of SIR1 protein establishes transcriptional silencing at HM loci and telomeres in yeast. Cell 75, 531–541.

(30) Rohner, S., Gasser, S. M., and Meister, P. (2008) Modules for cloning-free chromatin tagging in Saccharomyces cerevisae. Yeast 25, 235–239.

(31) Meinema, A. C., Laba, J. K., Hapsari, R. A., Otten, R., Mulder, F. A. A., Kralt, A., van den Bogaart, G., Lusk, C. P., Poolman, B., and Veenhoff, L. M. (2011) Long unfolded linkers facilitate membrane protein import through the nuclear pore complex. Science 333, 90–93.

(32) Egecioglu, D. E., D’Urso, A., Brickner, D. G., Light, W. H., and Brickner, J. H. (2014) Approaches to studying subnuclear organization and gene-nuclear pore interactions. Methods Cell Biol. 122, 463–485.

(33) Mali, P., Aach, J., Stranges, P. B., Esvelt, K. M., Moosburner, M., Kosuri, S., Yang, L., and Church, G. M. (2013) CAS9 transcriptional activators for target specificity screening and paired nickases for cooperative genome engineering. Nat. Biotechnol. 31, 833–838.

(34) Gilbert, L. A., Larson, M. H., Morsut, L., Liu, Z., Brar, G. A., Torres, S. E., Stern-Ginossar, N., Brandman, O., Whitehead, E. H., Doudna, J. A., Lim, W. A., Weissman, J. S., and Qi, L. S. (2013) CRISPR-mediated modular RNA-guided regulation of transcription in eukaryotes. Cell 154, 442–451.

(35) Zalatan, J. G., Lee, M. E., Almeida, R., Gilbert, L. A., Whitehead, E. H., La Russa, M., Tsai, J. C., Weissman, J. S., Dueber, J. E., Qi, L. S., and Lim, W. A. (2015) Engineering complex synthetic transcriptional programs with CRISPR RNA scaffolds. Cell 160, 339–350.

(36) Konermann, S., Brigham, M. D., Trevino, A. E., Joung, J., Abudayyeh, O. O., Barcena, C., Hsu, P. D., Habib, N., Gootenberg, J. S., Nishimasu, H., Nureki, O., and Zhang, F. (2015) Genome-scale transcriptional activation by an engineered CRISPR-Cas9 complex. Nature 517, 583–588.

(37) Ahmed, S., Brickner, D. G., Light, W. H., Cajigas, I., McDonough, M., Froyshteter, A. B., Volpe, T., and Brickner, J. H. (2010) DNA zip codes control an ancient mechanism for gene targeting to the nuclear periphery. Nat. Cell Biol. 12, 111–118.

(38) Brickner, D. G., Sood, V., Tutucci, E., Coukos, R., Viets, K., Singer, R. H., and Brickner, J. H. (2016) Subnuclear positioning and interchromosomal clustering of the GAL1-10 locus are controlled by separable, interdependent mechanisms. Mol. Biol. Cell 27, 2980–2993.

(39) Palladino, F., Laroche, T., Gilson, E., Axelrod, A., Pillus, L., and Gasser, S. M. (1993) SIR3 and SIR4 proteins are required for the positioning and integrity of yeast telomeres. Cell 75, 543– 555.

(40) Gotta, M., Laroche, T., Formenton, A., Maillet, L., Scherthan, H., and Gasser, S. M. (1996) The clustering of telomeres and colocalization with Rap1, Sir3, and Sir4 proteins in wild-type Saccharomyces cerevisiae. J. Cell Biol. 134, 1349–1363.

(41) Bystricky, K., Heun, P., Gehlen, L., Langowski, J., and Gasser, S. M. (2004) Long-range compaction and flexibility of interphase chromatin in budding yeast analyzed by high-resolution imaging techniques. Proc. Natl. Acad. Sci. USA 101, 16495–16500.

(42) Reece, R. J., and Ptashne, M. (1993) Determinants of binding-site specificity among yeast C6 zinc cluster proteins. Science 261, 909–911.

(43) Sternberg, S. H., Redding, S., Jinek, M., Greene, E. C., and Doudna, J. A. (2014) DNA interrogation by the CRISPR RNA-guided endonuclease Cas9. Nature 507, 62–67.

(44) Richardson, C. D., Ray, G. J., DeWitt, M. A., Curie, G. L., and Corn, J. E. (2016) Enhancing homology-directed genome editing by catalytically active and inactive CRISPR-Cas9 using asymmetric donor DNA. Nat. Biotechnol. 34, 339–344.

(45) Lim, F., and Peabody, D. S. (1994) Mutations that increase the affinity of a translational repressor for RNA. Nucleic Acids Res. 22, 3748–3752.

(46) Sharff, A. J., Rodseth, L. E., Spurlino, J. C., and Quiocho, F. A. (1992) Crystallographic evidence of a large ligand-induced hinge-twist motion between the two domains of the maltodextrin binding protein involved in active transport and chemotaxis. Biochemistry 31, 10657–10663.

(47) Lashkari, D. A., DeRisi, J. L., McCusker, J. H., Namath, A. F., Gentile, C., Hwang, S. Y., Brown, P. O., and Davis, R. W. (1997) Yeast microarrays for genome wide parallel genetic and gene expression analysis. Proc. Natl. Acad. Sci. USA 94, 13057–13062.

(48) Wu, X., Scott, D. A., Kriz, A. J., Chiu, A. C., Hsu, P. D., Dadon, D. B., Cheng, A. W., Trevino, A. E., Konermann, S., Chen, S., Jaenisch, R., Zhang, F., and Sharp, P. A. (2014) Genome-wide binding of the CRISPR endonuclease Cas9 in mammalian cells. Nat. Biotechnol. 32, 670–676.

(49) Horlbeck, M. A., Witkowsky, L. B., Guglielmi, B., Replogle, J. M., Gilbert, L. A., Villalta, J. E., Torigoe, S. E., Tjian, R., and Weissman, J. S. (2016) Nucleosomes impede Cas9 access to DNA in vivo and in vitro. eLife 5.

(50) Towbin, B. D., Gonzalez-Sandoval, A., and Gasser, S. M. (2013) Mechanisms of heterochromatin subnuclear localization. Trends in Biochemical Sciences 38, 356–363.

(51) Kumaran, R. I., and Spector, D. L. (2008) A genetic locus targeted to the nuclear periphery in living cells maintains its transcriptional competence. J. Cell Biol. 180, 51–65.

(52) Taddei, A., Van Houwe, G., Nagai, S., Erb, I., van Nimwegen, E., and Gasser, S. M. (2009) The functional importance of telomere clustering: global changes in gene expression result from SIR factor dispersion. Genome Res. 19, 611–625.

(53) Larson, A. G., and Narlikar, G. J. (2018) The role of phase separation in heterochromatin formation, function, and regulation. Biochemistry 57, 2540–2548.

(54) Zalatan, J. G. (2017) CRISPR-Cas RNA Scaffolds for Transcriptional Programming in Yeast. Methods Mol. Biol. 1632, 341–357.

(55) Meister, P., Gehlen, L. R., Varela, E., Kalck, V., and Gasser, S. M. (2010) Visualizing yeast chromosomes and nuclear architecture. Methods Enzymol 470, 535–567.

