## Supporting Information for "CRISPR-Cas-mediated tethering recruits the yeast *HMR* mating-type locus to the nuclear periphery but fails to silence gene expression"

This file contains:

- Supporting Figures 1-2
- Supporting Tables 1-3
- Guide RNA, effector protein, and reporter gene sequences
- Supporting References

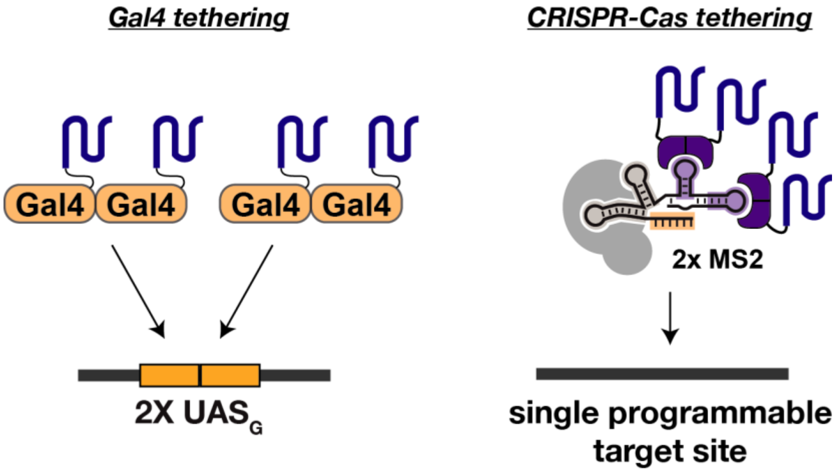

**Figure S1:** Gal4 and MCP are functional dimers. (A) The Gal4 DBD binds the UAS$_G$ site as a dimer.[1] A 2X UAS$_G$ site recruits four copies of Gal4-Yif1. (B) MCP binds to MS2 as a dimer.[2] A 2x MS2 scRNA that targets a single site recruits 4 copies of MCP-Yif1.

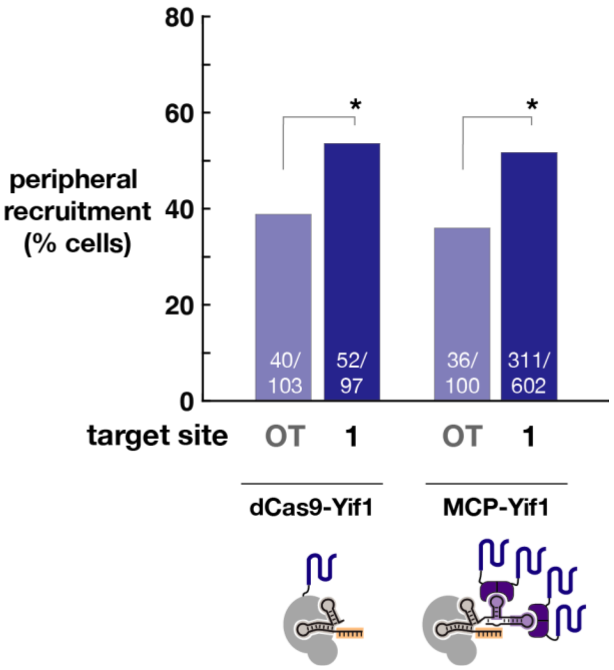

**Figure S2:** A direct dCas9-Yif1 fusion protein recruits the *HMR* locus to the periphery in yeast. With dCas9-Yif1 and a gRNA targeting *HMR* site 1 (Table S1), peripheral recruitment significantly increases compared to a strain with an off-target (OT) gRNA. Recruitment at the same target site mediated by a 2xMS2 scRNA and MCP-Yif1 (Fig 2C) is shown on the same plot for comparison. Peripheral recruitment was scored as described in the methods for yeast strains with and without recruitment systems. Exact values (recruited/total) are shown in white text with each bar. Statistical significance for a significant change in localization relative to a corresponding negative control was evaluated using a 2-tailed chi-squared test ($p$ value $\leq 0.05$, indicated by *). The $p$ value for dCas9-Yif1 recruitment is 0.05, and the $p$ value for MCP-Yif1 recruitment is $<0.0001$.

### Supplemental Tables

**Table S1.** gRNA target sites

| sgRNA target[a] | DNA Sequence | Location[b] |
|---|---|---|
| HMR_1 | ATTATATTGCAAAAACTCGA | ChrIII, -2496 (partially overlaps $UAS_G$) |
| HMR_2 | GGACAGTCCTCCGTCGACGG | ChrIII, -2472 (within $UAS_G$) |
| HMR_3 | TATTCTCCAAAACAATAATA | ChrIII, -8602 |
| HMR_4 | AATTCGAATAAGATAAACAG | ChrIII, -15223 |
| HMR_5 | AAAGGGTTTATATCCGAAGG | ChrIII, -102058 |
| HMR_6 | GGAGGACTGTCCTCCGTCGA | ChrIII, -2439 (within $UAS_G$) |
| HMR_7 | TCATGTACTAAACTAAAATC | ChrIII, -2414 |
| HMR_8 | AGAATAAGCGCAGGTACTCC | ChrIII, -2341 |
| W17 | GAAGTCAGTTGACAGAGTCG | Synthetic target site, used for off-target(OT) control in Fig 2 |
| GAL2_1 | CACATCACCAGACTTATCTC | ChrXII, +25 |
| GAL2_2 | TGAGAATGTTCGAACGATCC | ChrXII, +129 |
| GAL2_3 | AAATAAGTCAGGTACTTGCC | ChrXII, +61 |
| GAL2_4 | CGTCATTTAGGTCTAAAGTC | ChrXII, +382 |

[a] For *HMR* and *GAL2* target sites, see schematics in Figures 2 & 4 and complete reporter sequences below. For the HMR_6 site there are two repeats of the 20 base sequence in the $UAS_G$, but only one of these repeats has an appropriately positioned PAM.

[b] Location numbers given for *HMR* and *GAL2* sites are the distance from the 5' end (PAM distal) of the guide sequence to the integration sites for their respective tetO arrays.

**Table S2.** Yeast strains

| Strain | Description | Genotype |
|---|---|---|
| SO992 | W303 derivative | *MAT**a** ura3 leu2 trp1 his3 can1R ade* |
| yRK036 | *HMR* reporter | *SO992 Aeb:: 2x* $UAS_G$ *hmr::Trp1* |
| yRK045 | *HMR* reporter/dCas9/MCP-Yif1 | *SO992 Aeb:: 2x* $UAS_G$ *hmr::Trp1 Leu2::pTdh dCas9 His3::pAdh MCP-Yif1* |
| yRK113[a] | yRK036/tetO-array(*HMR*)/ tetR-GFP/mCherry-Heh2 | *SO992 Aeb:: 2x* $UAS_G$ *hmr::Trp1 TetO(60x)_LEU2 HO::pUra3 TetR-GFP_hph*$^R$ *mfa2::pTef1 mCherry-Heh2_KanMX* |
| yRK119[b] | yRK036/tetO-array(*HMR*)/ tetR-GFP/mCherry-Heh2 | *SO992 Aeb:: 2x* $UAS_G$ *hmr::Trp1 TetO HIS3 (60x) HO::pUra3 TetR-GFP_hph*$^R$ *mfa2::pTef1 mCherry-Heh2_KanMX* |
| yRK121 | yRK119/dCas9-Yif1 | *SO992 Aeb:: 2x* $UAS_G$ *hmr::Trp1 TetO HIS3 (60x) HO::pUra3 TetR-GFP_hph*$^R$ *mfa2::pTef1 mCherry-Heh2_KanMX Leu2::pTdh dCas9-Yif1* |
| yRK124 | yRK119/dCas9/MCP-Yif1 | *SO992 Aeb:: 2x* $UAS_G$ *hmr::Trp1 TetO HIS3 (60x) HO::pUra3 TetR-GFP_hph*$^R$ *mfa2::pTef1 mCherry-Heh2_KanMX Leu2::pTdh dCas9 pAdh MCP-Yif1* |
| yEC059[c] | mCherry-Heh2/tetR-GFP/tetO-array(*GAL2*)/dCas9/MCP-Yif1 | *S0992 mfa2::pTef1 mCherry Heh2 Kan HO::TetR-GFP_hph*$^R$ *Gal2::TetO(60x)_His3 LEU2::pTdh dCas9 pAdh MCP-Yif1* |

[a] yRK113 used pRS14 (Leu selection) to deliver the tetO array and is the parent for all Gal4-derivative strains in Fig 2.
[b] yRK119 used pRS8 (His selection) to deliver the tetO array and is the parent for all CRISPR-Cas tethering strains at *HMR* in Fig 2.
[c] yEC059 used pRS8 (His selection) to deliver the tetO array and is the parent strain for all *GAL2* recruitment strains in Fig 4.
pSR8 and pSR14 are described in Rohner et al., 2008.[3]

**Table S3.** Yeast protein expression plasmids

| **Plasmid**[a] | **Parent Vector**[b] | **Marker** | **Promoter** | **Gene** | **Terminator** |
|---|---|---|---|---|---|
| pJZC518 | pNH605 | *leu2* | *pTdh3* | dCas9 | *C. alb. Adh1* |
| pRK071 | pNH603 | *his3* | *pTdh3* | dCas9-$Yif1_{55\text{-}314}$ | *C. alb. Adh1* |
| pRK076 | pNH603 | *His3* | *pAdh* | MCP-$Yif1_{55\text{-}314}$ | *C. alb. Adh1* |
| pRK067 | p423 | *his3* | *pAdh* | $Gal4_{DBD}$-$Yif1_{55\text{-}314}$ | *cyc* |
| pEC091 | p423 | *his3* | *pAdh* | $Gal4_{DBD}$ | *cyc* |
| pRK144 | p423 | *his3* | *pAdh* | $Gal4_{DBD}$-MBP-$Yif1_{55\text{-}314}$ | *cyc* |
| pRK149 | pJW607 | *$hph^R$* | *pUra3* | TetR-GFP | *C. alb. Adh1* |
| pRK160 | pJW609 | *KanMX* | *pTef1* | mCherry-$Heh2_{138\text{-}378}$ | *C. alb. Adh1* |
| pRK159 | pNH605 | *leu2* | *1) pTdh3*<br>*2) pAdh* | 1) dCas9<br>2) MCP-$Yif1_{55\text{-}314}$ | *1) C. alb. Adh1*<br>*2) C. alb. Adh1* |

[a] pJZC518 for dCas9 expression and the general strategy for delivering MCP-effector proteins, either on separate integrating plasmids or together with dCas9 on a single integration cassette, have been described previously.[4]

[b] The pNH600 series of yeast single copy integration vectors has been described previously.[5]

### Guide RNA, Effector Protein, and *HMR* Reporter Sequences

*Guide RNA Sequence Designs*

sgRNA and scRNA sequences were constructed as described previously.[4] Alternative target sites were cloned with standard PCR methods.

Parent sgRNA

ATTATATTGCAAAAACTCGAGTTTTAGAGCTAGAAATAGCAAGTTAAAATAAGGCTAGTCCGTTATCA
ACTTGAAAAAGTGGCACCGAGTCGGTGGTGCTTTTTTTGTTTTTTATGTCT

2x MS2 scRNA

ATTATATTGCAAAAACTCGAGTTTTAGAGCTAGAAATAGCAAGTTAAAATAAGGCTAGTCCGTTATCA
ACTTGAAAAAGTGGCACCGAGTCGGTGCgggagcACATGAGGATCACCCATGTgccacgagcgACATG
AGGATCACCCATGTcgctcgtgttcccTTTTTTTGTTTTTTATGTCT

Annotations: 20 base target site (HMR_1), 2x MS2, SUP4 terminator

*Effector Protein Sequences*

>MCP$_{V29I\Delta FG}$-Yif1$_{55\text{-}314}$
MPKKKRKVGSMASNFTQFVLVDNGGTGDVTVAPSNFANGIAEWISSNSRSQAYKVTCSVRQSSAQNRKYTIKVEVPK
GAWRSYLNMELTIPIFATNSDCELIVKAMQGLLKDGNPIPSAIAANSGIYGSAGMGGFPFQDPRGSMAFQLGQSAFS
NFIGQDNFNQFQETVNKATANAAGSQQISTYFQVSTRYVINKLKLILVPFLNGTKNWQRIMDSGNFLPPRDDVNSPD
MYMPIMGLVTYILIWNTQQGLKGSFNPEDLYYKLSSTLAFVCLDLLILKLGLYLLIDSKIPSFSLVELLCYVGYKFV
PLILAQLLTNVTMPFNLNILIKFYLFIAFGVFLLRSVKFNLLSRSGAEDDDIHVSISKSTVKKCNYFLFVYGFIWQN
VLMWLMG

> Gal4$_{DBD}$-Yif1$_{55\text{-}314}$
MKLLSSIEQACDICRLKKLKCSKEKPKCAKCLKNNWECRYSPKTKRSPLTRAHLTEVESRLERLEQLFLLIFPREDL
DMILKMDSLQDIKALLTGLFVQDNVNKDAVTDRLASVETDMPLTLRQHRISATSSSEESSNKGQRQLTVSAAPEFGG
FPFQDPRGSMAFQLGQSAFSNFIGQDNFNQFQETVNKATANAAGSQQISTYFQVSTRYVINKLKLILVPFLNGTKNW
QRIMDSGNFLPPRDDVNSPDMYMPIMGLVTYILIWNTQQGLKGSFNPEDLYYKLSSTLAFVCLDLLILKLGLYLLID
SKIPSFSLVELLCYVGYKFVPLILAQLLTNVTMPFNLNILIKFYLFIAFGVFLLRSVKFNLLSRSGAEDDDIHVSIS
KSTVKKCNYFLFVYGFIWQNVLMWLMG

> Gal4$_{DBD}$-M-Yif1$_{55\text{-}314}$ (*M = maltose binding protein, MBP, 3x HA*)
MKLLSSIEQACDICRLKKLKCSKEKPKCAKCLKNNWECRYSPKTKRSPLTRAHLTEVESRLERLEQLFLLIFPREDL
DMILKMDSLQDIKALLTGLFVQDNVNKDAVTDRLASVETDMPLTLRQHRISATSSSEESSNKGQRQLTVSAAPEFMK
IEEGKLVIWINGDKGYNGLAEVGKKFEKDTGIKVTVEHPDKLEEKFPQVAATGDGPDIIFWAHDRFGGYAQSGLLAE
ITPDKAFQDKLYPFTWDAVRYNGKLIAYPIAVEALSLIYNKDLLPNPPKTWEEIPALDKELKAKGKSALMFNLQEPY
FTWPLIAADGGYAFKYENGKYDIKDVGVDNAGAKAGLTFLVDLIKNKHMNADTDYSIAEAAFNKGETAMTINGPWAW
SNIDTSKVNYGVTVLPTFKGQPSKPFVGVLSAGINAASPNKELAKEFLENYLLTDEGLEAVNKDKPLGAVALKSYEE
ELAKDPRIAATMENAQKGEIMPNIPQMSAFWYAVRTAVINAASGRQTVDEALKDAQTNSSSNNNNNNNNNNLGIEGR
ISTSQFGSGYPYDVPDYAGSGYPYDVPDYAGSGYPYDVPDYAGSGKFGGFPFQDPRGSMAFQLGQSAFSNFIGQDNF
NQFQETVNKATANAAGSQQISTYFQVSTRYVINKLKLILVPFLNGTKNWQRIMDSGNFLPPRDDVNSPDMYMPIMGL
VTYILIWNTQQGLKGSFNPEDLYYKLSSTLAFVCLDLLILKLGLYLLIDSKIPSFSLVELLCYVGYKFVPLILAQLL
TNVTMPFNLNILIKFYLFIAFGVFLLRSVKFNLLSRSGAEDDDIHVSISKSTVKKCNYFLFVYGFIWQNVLMWLMG

> dCas9-Yif1$_{55\text{-}314}$
MLEDKKYSIGLAIGTNSVGWAVITDEYKVPSKKFKVLGNTDRHSIKKNLIGALLFDSGETAEATRLKRTARRRYTRR
KNRICYLQEIFSNEMAKVDDSFFHRLEESFLVEEDKKHERHPIFGNIVDEVAYHEKYPTIYHLRKKLVDSTDKADLR
LIYLALAHMIKFRGHFLIEGDLNPDNSDVDKLFIQLVQTYNQLFEENPINASGVDAKAILSARLSKSRRLENLIAQL
PGEKKNGLFGNLIALSLGLTPNFKSNFDLAEDAKLQLSKDTYDDDLDNLLAQIGDQYADLFLAAKNLSDAILLSDIL
RVNTEITKAPLSASMIKRYDEHHQDLTLLKALVRQQLPEKYKEIFFDQSKNGYAGYIDGGASQEEFYKFIKPILEKM
DGTEELLVKLNREDLLRKQRTFDNGSIPHQIHLGELHAILRRQEDFYPFLKDNREKIEKILTFRIPYYVGPLARGNS
RFAWMTRKSEETITPWNFEEVVDKGASAQSFIERMTNFDKNLPNEKVLPKHSLLYEYFTVYNELTKVKYVTEGMRKP
AFLSGEQKKAIVDLLFKTNRKVTVKQLKEDYFKKIECFDSVEISGVEDRFNASLGTYHDLLKIIKDKDFLDNEENED
ILEDIVLTLTLFEDREMIEERLKTYAHLFDDKVMKQLKRRRYTGWGRLSRKLINGIRDKQSGKTILDFLKSDGFANR
NFMQLIHDDSLTFKEDIQKAQVSGQGDSLHEHIANLAGSPAIKKGILQTVKVVDELVKVMGRHKPENIVIEMARENQ
TTQKGQKNSRERMKRIEEGIKELGSQILKEHPVENTQLQNEKLYLYYLQNGRDMYVDQELDINRLSDYDVDAIVPQS
FLKDDSIDNKVLTRSDKNRGKSDNVPSEEVVKKMKNYWRQLLNAKLITQRKFDNLTKAERGGLSELDKAGFIKRQLV
ETRQITKHVAQILDSRMNTKYDENDKLIREVKVITLKSKLVSDFRKDFQFYKVREINNYHHAHDAYLNAVVGTALIK
KYPKLESEFVYGDYKVYDVRKMIAKSEQEIGKATAKYFFYSNIMNFFKTEITLANGEIRKRPLIETNGETGEIVWDK
GRDFATVRKVLSMPQVNIVKKTEVQTGGFSKESILPKRNSDKLIARKKDWDPKKYGGFDSPTVAYSVLVVAKVEKGK
SKKLKSVKELLGITIMERSSFEKNPIDFLEAKGYKEVKKDLIIKLPKYSLFELENGRKRMLASAGELQKGNELALPS
KYVNFLYLASHYEKLKGSPEDNEQKQLFVEQHKHYLDEIIEQISEFSKRVILADANLDKVLSAYNKHRDKPIREQAE
NIIHLFTLTNLGAPAAFKYFDTTIDRKRYTSTKEVLDATLIHQSITGLYETRIDLSQLGGDEASDPKKKRKVDPKKK
RKVDPKKKRKVAGSGGFPFQDPRGSMAFQLGQSAFSNFIGQDNFNQFQETVNKATANAAGSQQISTYFQVSTRYVIN
KLKLILVPFLNGTKNWQRIMDSGNFLPPRDDVNSPDMYMPIMGLVTYILIWNTQQGLKGSFNPEDLYYKLSSTLAFV
CLDLLILKLGLYLLIDSKIPSFSLVELLCYVGYKFVPLILAQLLTNVTMPFNLNILIKFYLFIAFGVFLLRSVKFNL
LSRSGAEDDDIHVSISKSTVKKCNYFLFVYGFIWQNVLMWLMG

*HMR reporter sequence*

The *HMR* reporter sequence (Fig 2A & S2) was designed following a previously described silencing reporter (*HMR* Aeb, with the E and B sites removed)[6,7] that contains a binding site for Gal4 (the UAS$_G$ or upstream activating sequence)[8] within the *HMR*-E region and a downstream Trp1 reporter gene integrated at the endogenous *HMR* locus in *S. cerevisiae* chromosome III. Genes at the *HMR* locus are normally silenced, but the UAS$_G$ insertion disrupts endogenous regulatory sites to allow gene expression. The Gal4$_{DBD}$ binds at the UAS$_G$, and Gal4$_{DBD}$ fusion proteins can rescue the silencing phenotype by directly recruiting silencing factors or by recruiting the *HMR* locus to the nuclear periphery.[6,7]

We integrated a tetO array approximately 2.4 kb downstream of the 2xUAS$_G$ site, which allows direct visualization of the locus with tetO-GFP. The original reporter described in the literature contains a 3X UAS$_G$ repeat,[6,7] while our reporter contains a 2X UAS$_G$ repeat. In functional silencing assays, the 2X UAS$_G$ reporter construct was effectively silenced by Gal4$_{DBD}$-Yif1 expression (Fig 2), similar to that described for the 3X UAS$_G$ reporter.[6]

Annotations[7,9,10]: *HMR*-E (Aeb), UAS$_G$, Trp1 (5'utr-ORF-utr-3'), *HMR*-I

```
ctagtacttaaaaaaactgtagtttcagtgcaaaaaagttttaacattacgtatcttgtacccttttttattgcatatagaaaggtc
aaataatccttcacatcatgaaatataagctaaatcgcatttcttttcgtccacatttgcaaacaaaacttttcaataatatttt
ataaatagtatcaatatatatatatatatttatttgtttactttttctatcagtgttttcaatttttttattaaacaatgtttgatt
ttttcaatcgcaatttaatacctaaatataaaaaatgttattatattgcaaaaaCTCGACGGAGGACAGTCCTCCGTCGACGGAGG
ACAGTCCTCCGTCGAGaatatttgaaagcaatagatcatgtactaaactaaaatcagggaaattaagactccttttgaagtaatac
ctattacttactaatacgtttgagaataagcgcaggtactcctggtttttgttaaaattacaaatttatacttagcattacgaaga
ttctcgattccgaaaaacaaaaattttatcgtcatatacaaatctagggtcgaaaaaagaaaaggagagggccaagagggagggca
ttggtgactattgagcacgtgagtatacgtgattaagcacacaaaggcagcttggagtATGTCTGTTATTAATTTCACAGGTAGTT
CTGGTCCATTGGTGAAAGTTTGCGGCTTGCAGAGCACAGAGGCCGCAGAATGTGCTCTAGATTCCGATGCTGACTTGCTGGGTATT
ATATGTGTGCCCAATAGAAAGAGAACAATTGACCCGGTTATTGCAAGGAAAATTTCAAGTCTTGTAAAAGCATATAAAAATAGTTC
AGGCACTCCGAAATACTTGGTTGGCGTGTTTCGTAATCAACCTAAGGAGGATGTTTTGGCTCTGGTCAATGATTACGGCATTGATA
TCGTCCAACTGCATGGAGATGAGTCGTGGCAAGAATACCAAGAGTTCCTCGGTTTGCCAGTTATTAAAAGACTCGTATTTCCAAAA
GACTGCAACATACTACTCAGTGCAGCTTCACAGAAACCTCATTCGTTTATTCCCTTGTTTGATTCAGAAGCAGGTGGGACAGGTGA
ACTTTTGGATTGGAACTCGATTTCTGACTGGGTTGGAAGGCAAGAGAGCCCCGAAAGCTTACATTTTATGTTAGCTGGTGGACTGA
CGCCAGAAAATGTTGGTGATGCGCTTAGATTAAATGGCGTTATTGGTGTTGATGTAAGCGGAGGTGTGGAGACAAATGGTGTAAAA
GACTCTAACAAAATAGCAAATTTCGTCAAAAATGCTAAGAAATAGgttattactgagtagtatttatttaagtattgtttgtgcac
ttgcctgcaggccttttgaaaagcaagcataaaaaataaattcgttttcaatgattaaaatagcatagtcgggtttttcttttagt
ttcagctttccgcaacagtataattttataaaccctggttttggttttgtagagtggttgacgaataattatgctgaagtacgtgg
tgacggatattgggaagatgtgtttgtacatttggccttatagagtgtggtcgtggcggaggttgtttatctttcgagtactgaat
gttgtcagtatagctatcctatttgaaactccccatcgtcttgctcttgttctcaatgtttgtttatatactcatatttctatgtg
tttatacaattgctattgtttatataatgtagtgacattttctcttaatcttatactaatttctatgacatttatataagaagaga
cttatgatcaacataattttgcaaactttgagagaaatatgtctttctactgcgataaagttattatttagattacatgtcaccaa
cattttcgtatatggcgatataaatttatcatgttttggtatgataatttaatttttaaaaaaacaaatttaattgacctcattaa
ttaatatttattaatacctttaatgttgaggtaaatagctattttctctcttcttttcctttagttggaatttgcacaagaaaatg
tttttccacacactttagcgtttttttcctaaatgttggaataaaaaacaactatcatctatcaaCTAGTAGTCACACTACCAATGT
GTTATCATTATACTGTGTTAAACAATGACATAAGGTATGAAAATTTGTCAACGAAGTTAGAGAAAGCTGGATGCAAGGATTGATAA
TGTGGTAGGAAAATGAAACATATAACGGAATGAGGAATAATCGTAATATCAGTATATAGAAATATAGATTCCCTTTTGAGGATTCC
TATATCCTCGAGGAGAACTTCTAGTATATTCTATATACCTAATATTATTACTTTTATCTACAATGCAACCCCACAATAATATAAAA
ATTCACCAATTCCGCATCTGCAGATTACTTTCCTAAATTTGCATATAGAATTGTCAAGCGCAAATCCGACGTCGATTCCGCGGCGG
ATGGGTCATTCTAGGTCATTCTACCAATTTTATTTGAGACCAGGTTTATTCAACCGGTAACATAGAAATATTCATACAATTAAGCT
TCTATGGCCAAGTTGGTAAGGCGCCACACTAGTAATGTGGAGATCATCGGTTCAAATCCGATTGGAAGCATTTTTTATCACGTTAT
TCGGTGACACCCAGGTTGCCGCCGCGTTCGCGTCCATCGTCATCTGAAAAATAATGAATATTAATGGACCTTGTGCCCCATAAAGG
TTCCATGTTCCATAAGTCTTCAATAATACTTTTGTATATTA[tetO_array_integration_site]ACCGTTATTCGGAGAT
CTCTTACGGCTTATGATTTTCTTTACATTCCAGGCCGCCTTTTG
```

*GAL2 sequence*

The *GAL2* (Chr XII) tetO array was designed with the array integrated 290 bases downstream of the Gal2 ORF, which allows direct visualization of the locus with tetO-GFP (Fig S2).

Annotation: Gal2 (ORF)

AATAGTAATAGTTAAGTAAACACAAGATTAACATAATAAAAAAAATAATTCTTTCATAATGGCAGTTGAGGAGAACAATATGCCTG
TTGTTTCACAGCAACCCCAAGCTGGTGAAGACGTGATCTCTTCACTCAGTAAAGATTCCCATTTAAGCGCACAATCTCAAAAGTAT
TCTAATGATGAATTGAAAGCCGGTGAGTCAGGGTCTGAAGGCTCCCAAAGTGTTCCTATAGAGATACCCAAGAAGCCCATGTCTGA
ATATGTTACCGTTTCCTTGCTTTGTTTGTGTGTTGCCTTCGGCGGCTTCATGTTTGGCTGGGATACCGGTACTATTTCTGGGTTTG
TTGTCCAAACAGACTTTTTGAGAAGGTTTGGTATGAAACATAAGGATGGTACCCACTATTTGTCAAACGTCAGAACAGGTTTAATC
GTCGCCATTTTCAATATTGGCTGTGCCTTTGGTGGTATTATACTTTCCAAAGGTGGAGATATGTATGGCCGTAAAAAGGGTCTTTC
GATTGTCGTCTCGGTTTATATAGTTGGTATTATCATTCAAATTGCCTCTATCAACAAGTGGTACCAATATTTCATTGGTAGAATCA
TATCTGGTTTGGGTGTCGGCGGCATCGCCGTCTTATGTCCTATGTTGATCTCTGAAATTGCTCCAAAGCACTTGAGAGGCACACTA
GTTTCTTGTTATCAGCTGATGATTACTGCAGGTATCTTTTTGGGCTACTGTACTAATTACGGTACAAAGAGCTATTCGAACTCAGT
TCAATGGAGAGTTCCATTAGGGCTATGTTTCGCTTGGTCATTATTTATGATTGGCGCTTTGACGTTAGTTCCTGAATCCCCACGTT
ATTTATGTGAGGTGAATAAGGTAGAAGACGCCAAGCGTTCCATTGCTAAGTCTAACAAGGTGTCACCAGAGGATCCTGCCGTCCAG
GCAGAGTTAGATCTGATCATGGCCGGTATAGAAGCTGAAAAACTGGCTGGCAATGCGTCCTGGGGGGAATTATTTTCCACCAAGAC
CAAAGTATTTCAACGTTTGTTGATGGGTGTGTTTGTTCAAATGTTCCAACAATTAACCGGTAACAATTATTTTTTCTACTACGGTA
CCGTTATTTTCAAGTCAGTTGGCCTGGATGATTCCTTTGAAACATCCATTGTCATTGGTGTAGTCAACTTTGCCTCCACTTTCTTT
AGTTTGTGGACTGTCGAAAACTTGGGACATCGTAAATGTTTACTTTTGGGCGCTGCCACTATGATGGCTTGTATGGTCATCTACGC
CTCTGTTGGTGTTACTAGATTATATCCTCACGGTAAAAGCCAGCCATCTTCTAAAGGTGCCGGTAACTGTATGATTGTCTTTACCT
GTTTTTATATTTTCTGTTATGCCACAACCTGGGCGCCAGTTGCCTGGGTCATCACAGCAGAATCATTCCCACTGAGAGTCAAGTCG
AAATGTATGGCGTTGGCCTCTGCTTCCAATTGGGTATGGGGGTTCTTGATTGCATTTTTCACCCCATTCATCACATCTGCCATTAA
CTTCTACTACGGTTATGTCTTCATGGGCTGTTTGGTTGCCATGTTTTTTTATGTCTTTTTCTTTGTTCCAGAAACTAAAGGCCTAT
CGTTAGAAGAAATTCAAGAATTATGGGAAGAAGGTGTTTTACCTTGGAAATCTGAAGGCTGGATTCCTTCATCCAGAAGAGGTAAT
AATTACGATTTAGAGGATTTACAACATGACGACAAACCGTGGTACAAGGCCATGCTAGAATAATGCGTTTGAAGTGAGACGCTCCA
TCATCTCTCTTAATTTTTCATGACTGACGTTTTTTCTTCATTTTAATTATCATAGTATTTGTTTGAAAAAAAAAAAAAAAAATTTC
CCTTATCAATGATATCCTTACGATTATATAAATTCCTTACCTAAACCTATTATTTGTGTACATATATCAGAGTATTATTACATATA
TAACCTTTTTCTCTAAAACAGGAAAAAAAAAAAGAAAACGATAACATGCTCTGCCATCCTTTGTTCACCGAGCAAAATTAAAAACGC
AAAATGAAT**[tetO_array_integration_site]**TGTCCCTATGAAATTATTAAAGGACCACATCACCAGACTTATCTCTGG
GGGGTCCTCTAGAAAATAAGTCAGGTACTTGCCTGGACTTTCTTCCAGTTGAATTCCTGAGCTAACATACAATTAATGGAGTGAGA
ATGTTCGAACGATCCAGGAGTTGCTTTTTTCAGTCATTTGTTTATCAGTGTAACAAGCATTTCCTTTATTTCTTTTATATCAACCT
GCAACCTATTAATATTCAATTCAACGTCTCGGAGCTTTTCTCGTACCGTTTTACCATCATCCATTTCCATGCTAACTAATTCTGAG
ATTACGATTACATATTGTCTCAGTTCTGGTTCATCAATGGCTGATAAAACTTTCATTACATTTTGGAAAAAATCTTCGTCATTTAG
GTCTAAAGTCAGGATTTGGTTCCTTATAGACTCTCTACGATCTTTTTTCAAGTTTTTTTCTGAAGAAATATGTCCATTTTCATTAG
TGACAGCGGTAGAGTCCTCTTCCGCTCGAACCAAACTATCACTGCCGGTGTTTTCCTCTCGAAAGGGTTTGTCTTTACCAATGGAA
AGACGCCCGTCTTCTGCGCTTTCATATAATTTGACAGTGGCTATATTGAACTGTTCTTGCAAATCGCCAGTTTTGGCATCCCCTAC
CCAACCTTTTTCATGCAATTCCTTTCCTTGAGTGACCC

### Supporting References

(1) Carey, M., Kakidani, H., Leatherwood, J., Mostashari, F., and Ptashne, M. (1989) An amino-terminal fragment of GAL4 binds DNA as a dimer. *J. Mol. Biol. 209*, 423–432.

(2) Valegård, K., Murray, J. B., Stockley, P. G., Stonehouse, N. J., and Liljas, L. (1994) Crystal structure of an RNA bacteriophage coat protein-operator complex. *Nature 371*, 623–626.

(3) Rohner, S., Gasser, S. M., and Meister, P. (2008) Modules for cloning-free chromatin tagging in *Saccharomyces cerevisae*. *Yeast 25*, 235–239.

(4) Zalatan, J. G., Lee, M. E., Almeida, R., Gilbert, L. A., Whitehead, E. H., La Russa, M., Tsai, J. C., Weissman, J. S., Dueber, J. E., Qi, L. S., and Lim, W. A. (2015) Engineering complex synthetic transcriptional programs with CRISPR RNA scaffolds. *Cell 160*, 339–350.

(5) Zalatan, J. G., Coyle, S. M., Rajan, S., Sidhu, S. S., and Lim, W. A. (2012) Conformational control of the Ste5 scaffold protein insulates against MAP kinase misactivation. *Science 337*, 1218–1222.

(6) Andrulis, E. D., Neiman, A. M., Zappulla, D. C., and Sternglanz, R. (1998) Perinuclear localization of chromatin facilitates transcriptional silencing. *Nature 394*, 592–595.

(7) Chien, C. T., Buck, S., Sternglanz, R., and Shore, D. (1993) Targeting of SIR1 protein establishes transcriptional silencing at HM loci and telomeres in yeast. *Cell 75*, 531–541.

(8) Bram, R. J., Lue, N. F., and Kornberg, R. D. (1986) A GAL family of upstream activating sequences in yeast: roles in both induction and repression of transcription. *EMBO J. 5*, 603–608.

(9) Abraham, J., Nasmyth, K. A., Strathern, J. N., Klar, A. J., and Hicks, J. B. (1984) Regulation of mating-type information in yeast. Negative control requiring sequences both 5′ and 3′ to the regulated region. *J. Mol. Biol. 176*, 307–331.

(10) Brand, A. H., Micklem, G., and Nasmyth, K. (1987) A yeast silencer contains sequences that can promote autonomous plasmid replication and transcriptional activation. *Cell 51*, 709–719.
